# Dengue virus strain 2 capsid protein switches the annealing pathway and reduces the intrinsic dynamics of the conserved 5’ untranslated region

**DOI:** 10.1101/2020.06.19.160978

**Authors:** Xin Ee Yong, Palur Venkata Raghuvamsi, Ganesh S. Anand, Thorsten Wohland, Kamal K. Sharma

## Abstract

The capsid protein of Dengue Virus strain 2 (DENV2C) is a structural protein with RNA chaperone activity that promotes multiple nucleic acid structural rearrangements, critical for transcription of the single-stranded positive-sense DENV2 genomic RNA. Annealing of the conserved 5’ untranslated region (5’UTR) to either its complementary sequence or to the 3’ untranslated region (3’UTR) occurs during (+)/(−) ds-RNA formation and (+) RNA circularization, respectively, both essential steps during DENV RNA replication. We investigated the effect of DENV2C on the annealing mechanism of two hairpin structures from the 5’UTR region (21-nt upstream AUG region (5’UAR) and 23-nt capsid-coding hairpin (5’cHP)) to their complementary sequences during (+)/(−) ds-RNA formation and (+) RNA circularization. Using fluorescence spectroscopy, DENV2C was found to switch annealing reactions nucleated mainly through kissing-loop intermediates to stem-stem interactions during (+)/(−) ds-RNA formation while it promotes annealing mainly through kissing-loop interactions during the (+) RNA circularization. Using FRET-FCS and trFRET, we determined that DENV2C exerts RNA chaperone activities by modulating intrinsic dynamics and by reducing the kinetically trapped unfavorable conformations of the 5’UTR sequence. Thus, DENV2C is likely to facilitate genome folding into functional conformations required for replication, playing a role in modulating (+)/(−) ds-RNA formation and (+) RNA circularization.

## INTRODCTION

Dengue fever is transmitted by the bite of an *Aedes aegypti* or *Aedes albopictus* mosquito carrying the dengue virus (DENV). DENV has four antigenically-distinct serotypes, DENV1-4, which complicate vaccine development because an effective vaccine should neutralize all four serotypes effectively (1). This is essential because secondary dengue infection tends to cause severe symptoms, such as dengue hemorrhagic fever and can even be fatal (2).

DENV2 is a positive-sense single-stranded RNA (ssRNA) virus with an icosahedral structure (T=3). The 50 nm virus has a genome of approximately 10.7 kb, which is translated to a polypeptide that is cleaved into three structural (envelope [E], pre-membrane [prM] and capsid [C]) and seven non-structural proteins (NS1, NS2A, NS2B, NS3, NS4A, NS4B, NS5) (3). The coding region is flanked at both ends by untranslated regions (UTR). The 5′ end has a type I cap structure (m^7^GpppAmp) mediating cap-dependent translation, but the virus can switch to a noncanonical translation mechanism when translation factors are limiting (4). Although the DENV 3’UTR lacks a poly(A) tail, the DENV genome is known to circularize, noted as (+) RNA circularization, to enhance its translation efficiency, similar to cellular mRNAs (Fig. 1A) (5, 6). This (+) RNA circularization of the DENV genome is proposed to proceed either through protein-nucleic acid interactions (6) or via long range RNA-RNA-based 5′ and 3′ (5′-3′) end interactions, which can occur in the absence of proteins (7–12). Thus far, two complementary sequences at the 5′ and 3′ ends have been identified for such long range 5’-3’ RNA end interactions and are necessary for the (+) RNA circularization (8). One of these sequences, a hairpin structure, known as the 5′ upstream AUG region (5′UAR) element in the 5′UTR, anneals with its complementary 3′UAR counterpart, which is located at the bottom part of 3′ stem loop (3’SL) in the 3’UTR (Fig. 1B) (7, 9, 12). In addition to the RNA sequences involved in 5′-3′-end interactions necessary for the (+) RNA circularization, the 5′ end of the viral genome harbours another functional RNA element capsid-coding region hairpin (5’cHP). The 5’cHP element resides within the capsid-coding region and directs start codon selection for RNA replication (13, 14). It is believed that the 5’cHP stalls the scanning initiation complex over the first AUG, favouring its recognition (15) and stabilizing the overall 5′-3′ panhandle structure (13). Apart from enhancing translation efficiency, (+) RNA circularization is also essential for the synthesis of the DENV (−) RNA genome, which serves as template for the amplification of additional positive strand genomic RNA (12). This (−) RNA genome synthesis includes NS5 binding at the 5′UTR and relocation of the polymerase at the 3′ initiation site via the (+) RNA circularization (16, 17). Thus, the hybridization of complementary sequences play a dual role: (i) to bring the NS5 polymerase-5’UTR promoter complex near the 3′ end of the genome and (ii) to open the large stem of the 3′SL structure by 5′-3′UAR annealing (18). As the (−) RNA genome is synthesized during transcription, secondary structures of the conserved 5’UTR RNA elements, like 5’UAR and 5’cHP, which is involved in the (−) RNA synthesis dissociates and form double-stranded (+)/(−) RNA (ds-RNA) to facilitate genome recombination (13). During this event, the 5’UAR and the 5’cHP dissociate and anneal to their complementary (−) RNA sequences, c5’UAR and c5’cHP, respectively (Fig. 1B).

**Figure 1:**
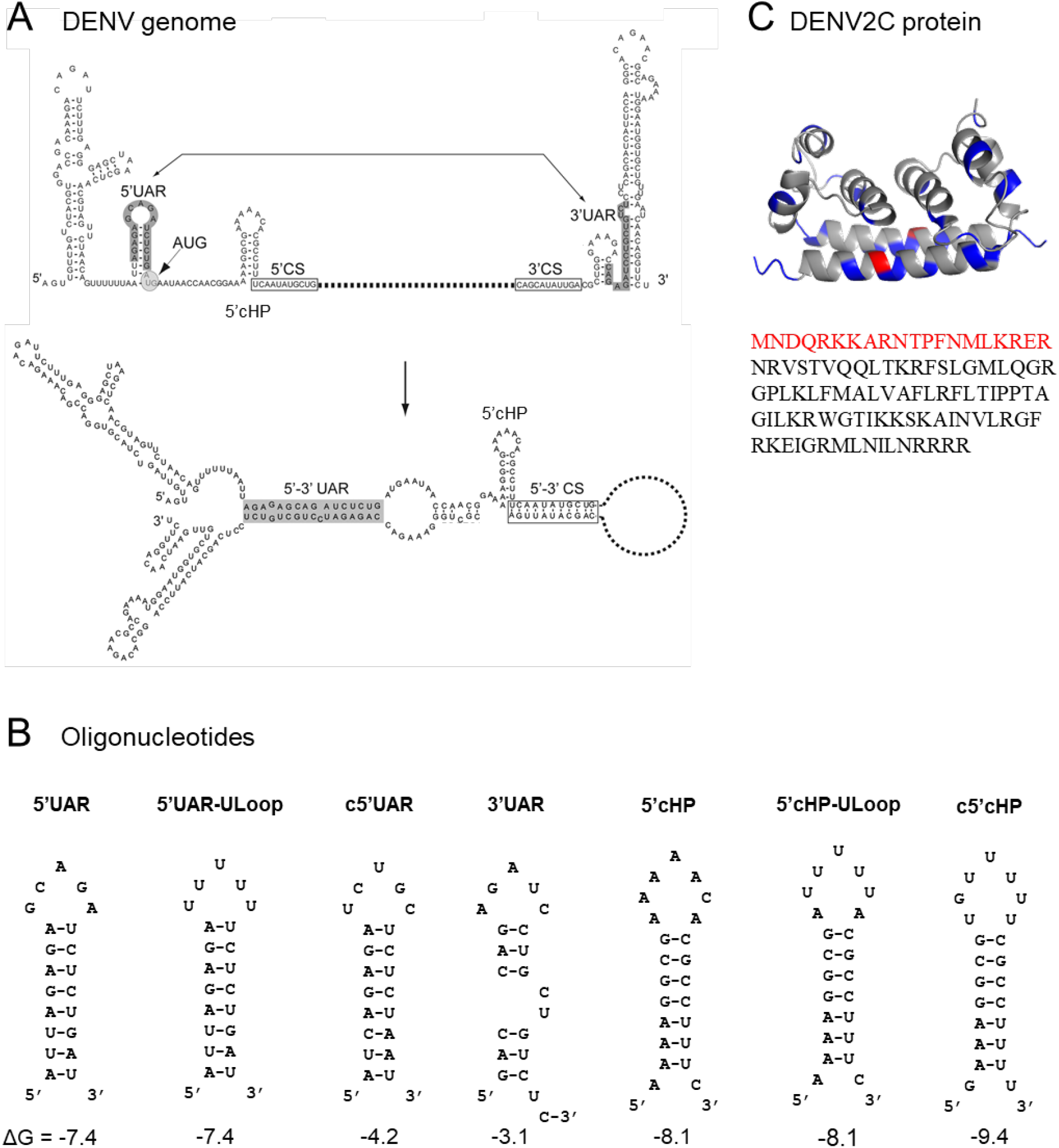
Structure of (A) DENV genome (adapted from (63)), (B) ODN sequences used in this study, and (C) DENV2C protein (PDB: 1R6R). (A) The 5’ and 3’ UTR region of DENV genome contain conserved circularization sequences, 5’-3’ UAR (upstream AUG region) (grey). (B) ODN sequences are derived from the 5’UAR, 5’cHP and 3’UAR region of the genome. Their secondary structures and ΔG values, shown below the structures in kcal/mol, were predicted using the mfold webtool (http://unafold.rna.albany.edu/). 5’UAR, 5’UAR-ULoop, 5’cHP and 5’cHP-ULoop are doubly labeled with 5’ FAM and 3’ TAMRA. Annealing pairs 5’UAR/c5’UAR and 5’cHP/c5’cHP represent (+)/(−) ds RNA, whereas 5’UAR/3’UAR are representing the (+) RNA circularization event. (C) The DENV2C protein is a homodimer. Blue and red residues highlight basic and acidic residues, respectively. In the protein sequence of DENV2C monomer, residues in red are part of the N-terminus disordered region which are not shown in the structure.

Furthermore, both (+)/(−) ds-RNA formation and the (+) RNA circularization require the aid of RNA chaperones. RNA chaperones help to prevent misfolding of RNA by allowing RNA to sample many different conformations (19). The DENV2 capsid protein, DENV2C, promotes annealing of the hammerhead ribozyme to its substrate, and promotes dissociation of the cleaved substrates, demonstrating RNA chaperoning ability (20). DENV2C exists as a homodimer, consisting of two 100-aa subunits. The 13 kDa DENV2C is highly basic, containing 26/100 basic amino acids and has a disordered region in the N-terminus, both key characteristics of many RNA chaperones (Fig. 1C) (21–23). The protein associates with the ssRNA genome to form the nucleocapsid and encapsulates the nucleocapsid in a lipid bilayer containing E and M proteins (24). DENV2C binds to RNA via its basic residues in helix-4 while binds to lipid bilayer using its hydrophobic cleft in helix-1 (12, 13). All further reference to DENV2C would be to one unit of the protein, i.e. the monomer.

Considering the abovementioned facts, DENV2C plays a vital role during virus formation possibly by modulating the interplay of RNA elements in the virus genome. Although, the interaction of RNA elements in the DENV 5’ and 3’ ends is required for viral RNA replication, the role of DENV2C during this interplay is still not explored. We hypothesize that, due to its chaperone properties, the DENV2C probably plays a role in modulating (+)/(−) ds-RNA formation and the (+) RNA circularization.

To achieve this aim, we characterized the role of DENV2C as an RNA chaperone by investigating the annealing kinetics of 5’UAR and 5’cHP to their complementary sequences during (+)/(−) ds-RNA formation and the (+) RNA circularization. To represent the (−) RNA synthesis or (+)/(−) ds-RNA formation, we investigated annealing kinetics of the 21-nt long 5’UAR and 23-nt long 5’cHP with their complementary c5’UAR and c5’cHP, respectively (Fig. 1B). On the other hand, the (+) RNA circularization event is represented by annealing kinetics of the 21-nt long 5’UAR with its complementary 3′UAR counterpart (Fig. 1B), which is located at the bottom part of 3′SL of the viral RNA. We doubly labeled 5’UAR and 5’cHP with 6-carboxyfluorescein (FAM) as donor fluorophore at the 5’ end and carboxytetramethylrhodamine (TAMRA) as acceptor fluorophore at the 3’ end, forming a Förster resonance energy transfer (FRET)-pair. Due to the proximity of donor and acceptor dyes in hairpin conformation of both 5’UAR and 5’cHP, the fluorescence of FAM is quenched. With the addition of a complementary sequence, the hairpin converts into a double-stranded sequence (extended duplex), in turn increasing the distance between donor and acceptor fluorophores. This increase in distance leads to florescence recovery of the donor that will provide information about the real-time annealing kinetics. By monitoring the annealing kinetics of the native and mutated 5’UAR and 5’cHP, we found that DENV2C promotes all three 5’UAR/c5’UAR, 5’cHP/c5’cHP and 5’UAR/3’UAR annealing. Interestingly, DENV2C is also able to switch the 5’UAR/c5’UAR annealing pathway that predominantly nucleates via the hairpin loops to a reaction pathway, nucleating through the hairpin stems. However, DENV2C does not alter the 5’UAR/3’UAR annealing pathway. We also determined by using FRET-fluorescence correlation spectroscopy (FRET-FCS) and time-resolve FRET (trFRET) that DENV2C exerts its chaperone functioning by reducing intrinsic dynamics of the 5’UAR and probably favoring one of the active conformations of the RNA hairpin. We also proposed mechanisms for the 5’UAR/c5’UAR and the 5’UAR/3’UAR annealing and compared the effects of DENV2C on (+)/(−) ds-RNA formation and the (+) RNA circularization. Our results suggest that DENV2C probably modulates (+)/(−) ds-RNA formation and the (+) RNA circularization events by modulating their annealing pathways.

## MATERIALS AND METHODS

### Oligonucleotides

All oligonucleotides (ODNs) were synthesized by Integrated DNA Technologies (Singapore). The doubly labeled ODNs (5’UAR, 5’UAR-ULoop, 5’cHP and 5’cHP-ULoop) were synthesized with 6-carboxyfluorescein (FAM) at the 5’ end and carboxytetramethylrhodamine (TAM) at the 3’ end. All ODNs were purified by the manufacturer using HPLC.

### DENV2C protein synthesis and purification

pET-21a(+) vector containing DENV2C protein gene sequence with N-terminal His tag and Tobacco Etch Virus (TEV) digestion site was purchased from GenScript (China). Recombinant capsid protein from DENV2 NGC strain was expressed in *Escherichia coli* BL21 strain. Transformed cells were grown in Luria-Bertani (LB) media supplemented with ampicillin (100 μg/ml) at 37°C until an optical density, OD_600_ of 0.6-0.8 was reached. A final concentration of 1 mM of isopropyl β-D-1-thiogalactopyranoside (IPTG) was used to induce expression and incubated for 16 to 18 hours at 18°C. This overnight culture was harvested by centrifugation at 6000 × g for 15 minutes at 4°C. The cells were lysed by sonication in NTE buffer consisting of 50mM Tris-HCl, 1 M NaCl, pH 7.5 with 0.8 mM DTT, protease inhibitor cocktail (COEDTAF-RO, Merck) and 40 μg/ml of RNase A (Thermo Fisher Scientific). The cell pellet was separated by centrifugation at 13000 rpm for 30 minutes, and the supernatant was collected and incubated with Cobalt beads for affinity chromatography. Capsid protein was eluted at 1 M imidazole in NTE buffer and the collected elute was subjected to size exclusion chromatography (HiLoad 16/600 Superdex 200, GE Healthcare). The fraction containing DENV2C protein were pooled and dialyzed against 50 mM HEPES, 150 mM NaCl, pH 7.5 buffer to remove imidazole and reduce NaCl concentration. Dialyzed DENV2C protein was concentrated using Amicon Ultra-4 Centrifugal Filter Units (10 kDa molecular weight cut off, Millipore). The concentration of the purified protein sample was determined using nanodrop.

### Fluorescence spectroscopy

Fluorescence spectroscopy measurements were done on Cary Eclipse Fluorescence Spectrophotometer (Agilent) with a temperature control module, using Hellma® fluorescence cuvettes with an internal chamber volume of 45 μL. Excitation and emission wavelengths of 480 nm and 520 nm were used to track the intensity of 5’ FAM on doubly labeled ODNs in real-time. Reactions were done in pseudo first order conditions, with the concentration of non-labeled ODN being at least 10-fold more than the doubly labeled ODN. Equal volumes of both reactants were mixed at the start of the reaction to prevent high local concentrations of either reactant. For reactions in the presence of DENV2C, the protein was added to each ODN and mixed well before the two reactants were mixed. DENV2C:ODN ratios of 2:1 was used for all reactions in the presence of DENV2C protein. All reactions were done in 50 mM HEPES, 30 mM NaCl, 0.2 mM MgCl_2_, pH 7.5 buffer and at 20°C, unless otherwise stated. Temperature-dependence experiments to obtain Arrhenius parameters were done by equilibrating both reactants at the specified temperature for 10 minutes prior to mixing. All curve fitting was done on OriginProTM software (ver 9.55).

### FCS and FRET-FCS

FCS measurements were carried out on a commercial Olympus FV1200 laser scanning confocal microscope (Olympus, Singapore) equipped with an FCS upgrade kit (PicoQuant, Berlin, Germany). Doubly labeled ODN samples were excited with a 543 nm continuous wave laser. The beam was focused onto the sample by a water immersion objective (60×, NA 1.2; Olympus, Singapore) after being reflected by a dichroic mirror (DM405/485/543/635, Olympus, Singapore) and the scanning unit. The 3’TAMRA fluorescence from 5’UAR was recorded by a single molecule avalanche photodiode (SPAD) (SPCM-AQR-14, PerkinElmer Optoelectronics, Quebec, Canada), through a 600/50 band pass emission filter. Detected photon counts are registered by a TimeHarp 260 time-correlated single photon counting board (PicoQuant). Autocorrelation analysis was done using SymPhoTime 64 (PicoQuant, Berlin, Germany) to obtain diffusion time and number of fluorescent particles. ODN and DENV2C protein mixtures were incubated for 10 minutes before measurements. Measurements were done at 25°C.

For FRET-FCS, the fluorescence signal from doubly-labeled 5’UAR was spectrally divided into donor and acceptor channels by a 560 dichroic longpass (DCLP) mirror. The donor and acceptor fluorescence were recorded using a set of single-molecule avalanche photodiodes (SPADs) (SPCM-AQR-14, PerkinElmer Optoelectronics, Quebec, Canada), through a 513/17 and 615/45 band pass emission filter (Omega, VT), respectively. The intensities of the donor and acceptor channel are collated in 20 μs time bins in the SymPhoTime64 software and exported. Using a home-written MATLAB script, the proximity ratio was calculated for each time bin and the autocorrelation of the proximity ratio was calculated. The calculated autocorrelation of the proximity ratio was then fitted using the Levenberg-Marquardt iteration algorithm by Origin 9.1.

### Time Resolved FRET (trFRET)

trFRET measurements were carried out on the same commercial Olympus FV1200 laser scanning confocal microscope equipped with a time-resolved LSM upgrade kit (Microtime 200, PicoQuant, GmbH, Berlin, Germany). Doubly labeled 5’UAR ODN was excited with a 485 nm pulsed diode laser with a 20 MHz repetition rate and 29 mW power (PDL series, Sepia II combiner module). The beam was focused into the sample by a water immersion objective (60×, NA 1.2; Olympus, Singapore) after being reflected by a dichroic mirror (DM405/485/543/635, Olympus, Singapore) and the scanning unit. The fluorescence was collected by the same objective followed by a pinhole (120 mm) to remove out-of-focus light. The fluorescence signal was spectrally divided into donor (green) and acceptor (red) channels by a 560 DCLP mirror. The 5’FAM donor fluorescence was recorded by a SPAD (SPCM-AQR-14, PerkinElmer Optoelectronics, Quebec, Canada), through a 513/17 band pass emission filter (Omega, VT). This donor signal was further processed by a time correlated single photon counting card (TimeHarp 260, PicoQuant) to build up the histogram of photon arrival times. The trFRET measurements were recorded for 180 s after incubating 5’UAR and DENV2C samples for 10 min. The mean lifetime (τ) was calculated from the individual fluorescence lifetimes (τ_i_) and their relative amplitudes (a_i_) according to (*τ*) = ∑ *α*_*i*_*τ*_*i*_. Donor fluorescence lifetime decay data were treated using the software SymPhoTime 64 (PicoQuant, GmbH). In all cases, the *χ*^2^ values were close to 1 and the weighted residuals as well as their autocorrelation were distributed randomly around 0, indicating a good fit. The reported values are mean and S.D.’s from at least three replicates.

## RESULTS

### The 5’UAR/c5’UAR and 5’cHP/c5’cHP annealing is faster as compared to the 5’UAR/3’UAR annealing

Annealing kinetics of doubly labeled 5’UAR to its non-labeled c5’UAR were monitored by tracking the increase in FAM fluorescence (Fig. 2A). Comparing the emission spectra of doubly labeled 5’UAR at the start and end of the annealing reaction shows a decrease in FRET with the formation of the extended duplex (Fig. 3A) as seen by an increase in FAM fluorescence and a corresponding decrease in TAMRA fluorescence. The real-time fluorescence intensity traces of the 5’UAR/c5’UAR reaction kinetics were fitted to a biexponential equation [1], where *I(t)* is the actual fluorescence intensity at 520 nm, upon excitation at 480 nm, *k*_obs1_ and *k*_obs2_ are the pseudo first order fast and slow reaction rates, *a* is the relative amplitude of the fast component and *t*_0_ is the start time of the reaction. *I_0_* and *I_f_* is the fluorescence intensity of doubly-labeled 5’UAR in the free state and in the final extended duplex (ED), respectively.

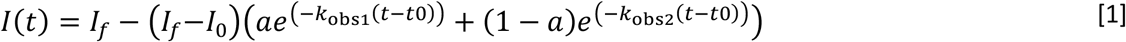

**Figure 2:**
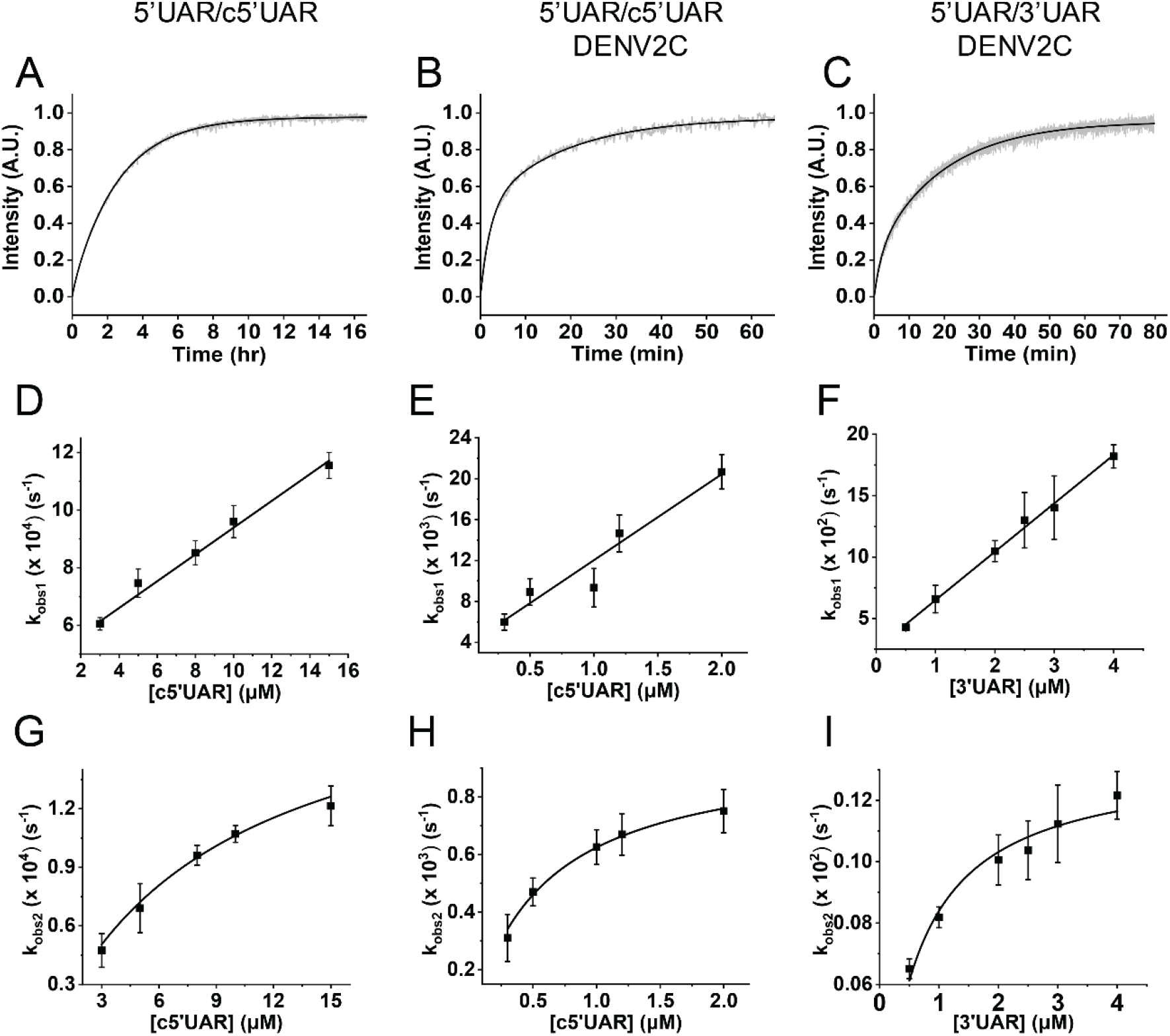
Real-time progress curves (A-C) and kinetic parameters (D-I) of annealing between 5’UAR and its complementary sequences (c5’UAR and 3’UAR) in the presence and absence of DENV2C. (A-C) Real-time progress curves (grey) and their fits to Equation [1] (black) to obtain fast (*k*_obs1_) and slow (*k*_obs2_) kinetic parameters. Real-time progress curve of the annealing between (A) 10 nM 5’UAR and 8 μM c5’UAR (B) 10 nM 5’UAR and 1 μM c5’UAR in the presence of DENV2C with DENV2C:ODN ratio of 2:1 and (C) 10 μM 5’UAR and 3 μM 3’UAR in the presence of DENV2C with DENV2C:ODN ratio of 2:1. The obtained values of *k*_obs1_ and *k*_obs2_ for (D and G) 5’UAR/c5’UAR, (E and H) 5’UAR/c5’UAR in presence of DENV2C and (F and I) 5’UAR/3’UAR in presence of DENV2C were plotted against corresponding complementary ODNs and fitted to Equation [2] and [3], respectively. Excitation and emission wavelengths used were 480 nm and 520 nm, respectively. Error bars show standard deviation from at least three repeats.

**Figure 3:**
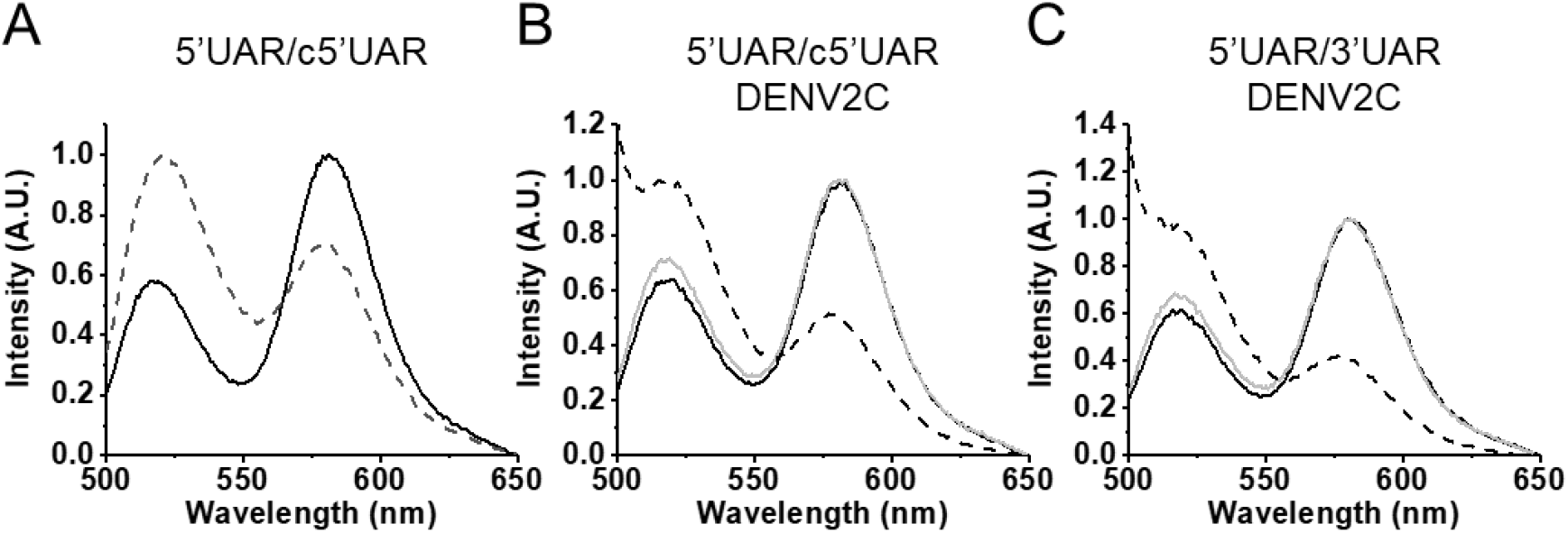
Emission spectra of 5’UAR before (solid black) and after annealing (dotted black) to its complementary sequences. (A) Emission spectra before and after annealing of 10 nM 5’UAR to 8 μM c5’UAR. (B) Emission spectra of 5’UAR in the absence (solid black) and presence (solid grey) of DENV2C in an DENV2C:ODN ratio of 2:1. The emission spectra after annealing of 10 nM 5’UAR to 1 μM c5’UAR in the presence of DENV2C is shown in dotted lines. (C) Emission spectra of 5’UAR in the absence (solid black) and presence (solid grey) of DENV2C in an DENV2C:ODN ratio of 2:1. The emission spectra after annealing of 10 nM 5’UAR to 3 μM 3’UAR in the presence of DENV2C is shown in dotted lines. The excitation wavelength was 480 nm.

The fast and slow reaction rates, *k*_obs1_ and *k*_obs2_, were plotted against the concentration of non-labeled reactant, c5’UAR (Fig. 2D and Fig. 2G). The fast component, *k*_obs1_, linearly varies with increasing c5’UAR concentrations ([c5’UAR]), while the slow component, *k*_obs2_, shows a hyperbolic dependence. The linear relationship between *k*_obs1_ and [c5’UAR] follows Equation [2] (27).

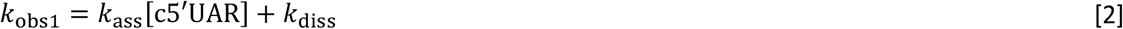

The hyperbolic dependence of *k*_obs2_ on increasing [c5’UAR] can be described in Equation [3], where K_a_ is the equilibrium constant governing IC formation, *k*_f_ and *k*_b_ are forward and backward interconversion kinetic rate constants.

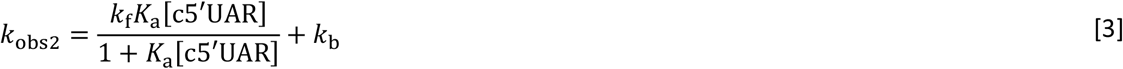

Fitting the linear and hyperbolic plots of *k*_obs1_ and *k*_obs2_ against increasing [c5’UAR] with Equations [2] and [3], respectively, generated kinetic parameters shown in Table 1.

**Table 1:**
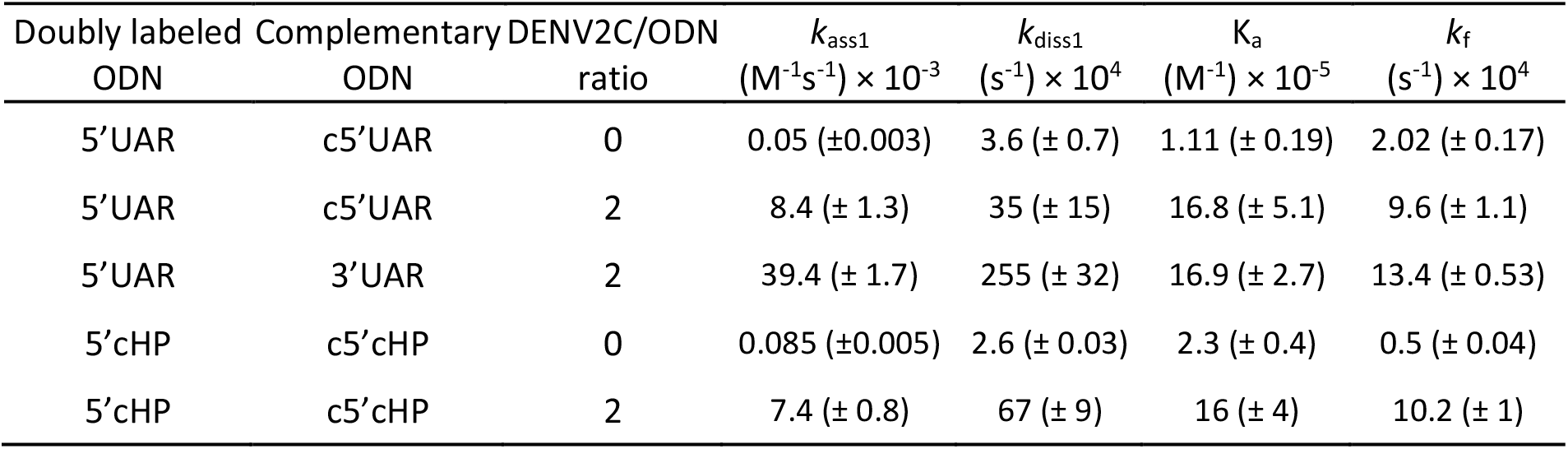
Kinetic parameters of 5’UAR/c5’UAR, 5’cHP/c5’cHP and 5’UAR/3’UAR annealing and their mutants in the absence and presence of DENV2C. Kinetic rate constants were calculated from the dependence of the *k*_obs_ values on the concentration of the unlabeled ODN, as indicated in Fig. 1. The *k*_ass_ and *k*_diss_ values were calculated with Equation [2], while the K_a_ and *k*_f_ values were calculated using Equation [3]. The K_a_ values were found to differ by a factor of <1.5 from the *k*_ass_/*k*_diss_ values, which further supports the proposed reaction Scheme 1 and 2.

Based on our acquired kinetic parameters, a reaction mechanism with single kinetic pathway starting from a 5’UAR species can be proposed:

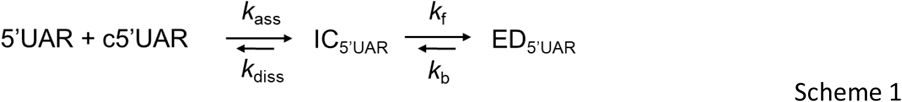

where a fast pre-equilibrium intermediate complex, IC_5’UAR_, precedes the formation of the final stable extended duplex, ED_5’UAR_, through a monomolecular reaction (28). The formation of IC_5’UAR_ is governed by the second order association constant, *k*_ass_, and the first order dissociation constant, *k*_diss_, whereas the formation of ED_5’UAR_ is governed by the forward and backward interconversion kinetic rate constants, *k*_f_ and *k*_b_ respectively. The hyperbolic dependence of *k*_obs2_ with increasing [c5’UAR] can be attributed to IC_5’UAR_ accumulation because of its slow interconversion to ED_5’UAR_. This is likely the rate-limiting step of the 5’UAR/c5’UAR annealing.

The *k*_ass_ value of 50 (± 3) M^−1^s^−1^ for the 5’UAR/c5’UAR annealing is at least 4 orders of magnitude smaller than the rate constants reported for annealing of unstructured sequences (10^5^-10^7^ M^−1^s^−1^), suggesting that there is a low probability for the reaction to occur at room temperature (29). The *k*_diss_ and *k*_b_ values were 3.5 (± 1.5) × 10^−3^ s^−1^ and close to zero, respectively, suggesting that dissociation of both IC_5’UAR_ and ED_5’UAR_ is negligible (Table 1).

To validate the postulated annealing mechanism (30–33), we used the Dynafit numerical resolution software (34), which allows simultaneous fitting of the experimental progress curves obtained at different [c5’UAR] (Fig. S1). The best estimates of the elementary rate constants *k*_ass_, *k*_diss_ and *k*_f_ (Table S1) were in excellent agreement with those found by the empirical approach (Table 1).

Further insights into the nature of the 5’UAR/c5’UAR annealing pathway were obtained from the temperature dependence of *k*_obs_ values, using the Arrhenius equation [4], which can be rewritten in a linear form [5], where the rate constant *k* is given by *k*_obs_/[c5’UAR], *A* is the pre-exponential Arrhenius factor, *E*_a_ is the activation energy, *R* is the universal gas constant and *T* is the temperature (in Kelvin).

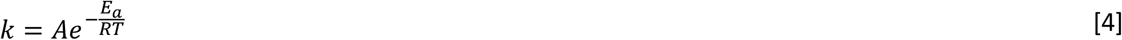

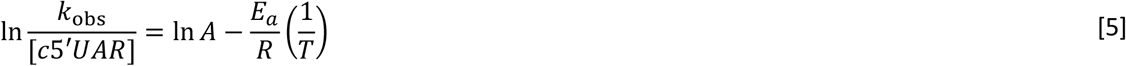

We observed an increase in both reaction rates leading to positive enthalpy values of the transition states of 11.9 (± 0.4) kcal/mol and 18.0 (± 1.0) kcal/mol for the fast and the slow components, respectively (Fig. 4 and Table 2). This suggests a pre-melting of 2-3 base-pairs (bp) of hydrogen bonds in 5’UAR hairpin structure (35, 36).

**Figure 4:**
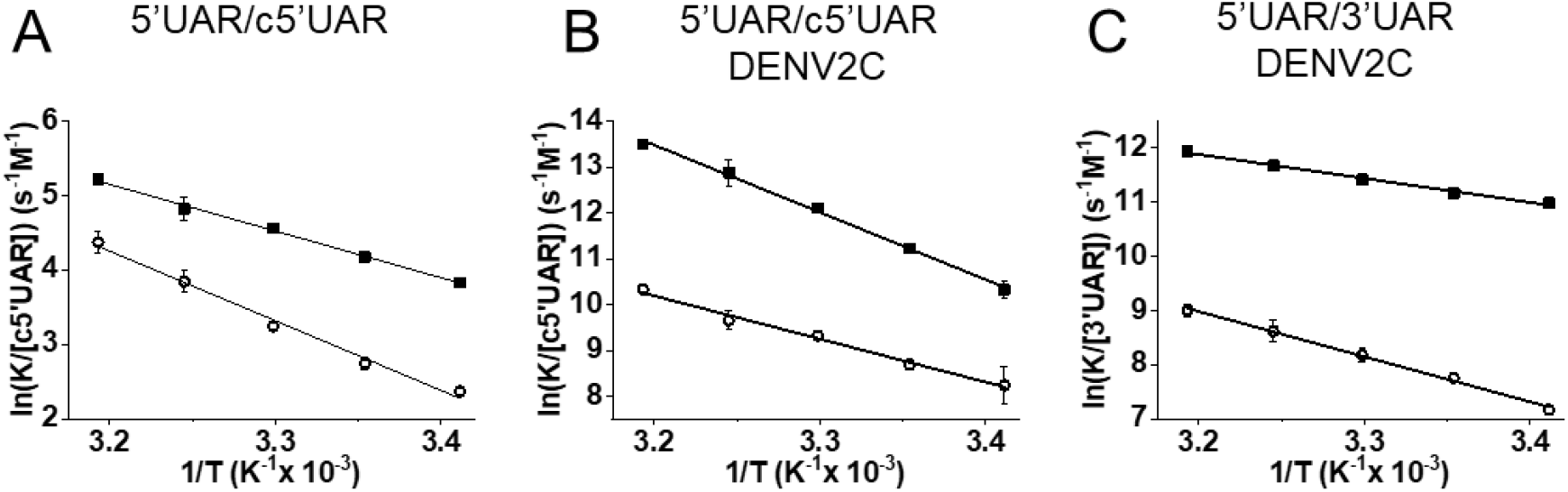
Temperature dependence of 5’UAR/c5’UAR and 5’UAR/3’UAR annealing in the (A) absence and (B and C) presence of DENV2C. The reactions between (A) 10 nM 5’UAR and 10 μM c5’UAR, (B) 10 nM 5’UAR and 500 nM c5’UAR in the presence of DENV2C and (C) 10 nM 5’UAR and 500 nM 3’UAR in the presence of DENV2C were monitored at various temperatures (20°C, 25°C, 30°C, 35°C and 40°C) to obtain fast, *k*_obs1_ and slow, *k*_obs2_ reaction rates. Natural logarithms of the fast (solid squares) and slow (open circles) reaction rates were plotted against the inverse of temperature. Solid lines are fits to Equation [4]. The Ea obtained from fitting were used to calculate transition state enthalpies (Table 2) using Δ*H* = *E_a_* − RT with T = 293.15 K. Excitation and emission wavelengths were 480 nm and 520 nm, respectively. Error bars represents the standard deviations of at least three repeats.

**Table 2:** Arrhenius parameters of 5’UAR/c5’UAR, 5’cHP/c5’cHP and 5’UAR/3’UAR annealing and their mutants in the absence and presence of DENV2C. The values of the transition state enthalpies, ΔH for the fast and slow pathways were calculated from the fits of temperature dependence of reaction rates, *k*_obs_, values to Equation [5], as described in Fig. 4.

| Doubly labeled ODN | Complementary ODN | DENV2C/ODN ratio | $\Delta H$ (Fast)<br>(kcal/mol) | $\Delta H$ (Slow)<br>(kcal/mol) |
| --- | --- | --- | --- | --- |
| 5'UAR | c5'UAR | 0 | 11.9 ± 0.4 | 18.0 ± 1.0 |
| 5'UAR | c5'UAR | 2 | 28.6 ± 0.9 | 18.1 ± 1.1 |
| 5'UAR | 3'UAR | 2 | 8.2 ± 0.4 | 16.0 ± 0.6 |

Next, similar to the 5’UAR/c5’UAR annealing, the 5’cHP/c5’cHP annealing shows a linear and hyperbolic dependence of *k*_obs1_ and *k*_obs2_ values against increasing [c5’cHP] (Fig. S2), thus suggesting a similar reaction mechanism as to that proposed by Scheme 1 (shown by Scheme a in the supplementary). The formation of IC_cHP_ was found to be ~1.7-fold faster in the 5’cHP/c5’cHP annealing compared to the 5’UAR/c5’UAR annealing with *k*_ass_ values of 85 (±5) M^−1^s^−1^ and 50 (± 3) M^−1^s^−1^, respectively (Table 1). However, the interconversion from IC_cHP_ to ED_cHP_ is slower by ~4-fold for 5’cHP/c5’cHP annealing compared to that of 5’UAR/c5’UAR annealing (Table 1), which can be attributed to higher stability of the 5’cHP hairpin, with ΔG_cHP_ of −8.1 kcal/mol compared to ΔG_5’UAR_ of −7.4 kcal/mol (Fig. 1).

As mentioned earlier, investigating the 5’UAR/3’UAR annealing will provide insights about the (+) RNA circularization. Interestingly, the 5’UAR/3’UAR annealing was drastically slower as compared to both 5’UAR/c5’UAR and 5’cHP/c5’cHP annealing and no increase in donor emission was observed even after 12 hours (Fig. 2A). This indicates that the kinetic barrier during the extended duplex (ED_3’UAR_) formation for the 5’UAR/3’UAR annealing is considerably higher as compared to both 5’UAR/c5’UAR and 5’cHP/c5’cHP annealing. This higher kinetic barrier also points towards the partial non-complementarity between 5’UAR and 3’UAR hairpins during the 5’UAR/3’UAR annealing (Fig. S3B).

Due to the extremely slow reaction kinetics of the 5’UAR/3’UAR annealing, no kinetic parameters could be determined, at least under our experimental conditions (Fig. S3E).

### DENV2C accelerates all three 5’UAR/c5’UAR, 5’cHP/c5’cHP and 5’UAR/3’UAR annealing

To understand the RNA chaperone activity of DENV2C and its role in genome recombination, we characterized the annealing of 5’UAR/c5’UAR, 5’cHP/c5’cHP and 5’UAR/3’UAR in the presence of DENV2C. However, RNA chaperones are known to cause nucleic acid aggregation (30, 37–40) which can cause adverse bias when using fluorescence-based techniques (37, 41). Thus, we first determined non-aggregating experimental conditions. Nucleic acid aggregation by positively charged proteins is concentration dependent (29, 42) and can be investigated using FCS (30, 40). In FCS, the fluorescence intensity arising from confocal volume (about 0.2 fL) is correlated to obtain information about the processes that give rise to fluorescence fluctuations. These fluctuations are governed by the diffusion of fluorescent species in the confocal volume, and thus parameters like average number of fluorescent species and their diffusion constant can be determined. Aggregation of labeled 5’UAR molecules by the DENV2C would be expected to decrease the number of fluorescent ODNs (N) and an increase in the diffusion time (τ_D_). By adding increasing concentrations of DENV2C to labeled ODN, we found no change in N or τ_D_ up to a DENV2C/ODN molar ratio of 2:1, indicating that no aggregation occurred under these conditions (Fig. S4). Thus, we selected a DENV2C/ODN molar ratio of 2:1 to characterize the annealing of 5’UAR/c5’UAR, 5’cHP/c5’cHP and 5’UAR/3’UAR in the presence of DENV2C.

Addition of DENV2C to the labeled 5’UAR or 5’cHP sequences did not lead to any significant change in their fluorescence spectrum (compare black and grey emission spectra in Fig. 3B and 3C), indicating that DENV2C was unable to destabilize the stem of their secondary structures, similar to other RNA chaperones (30, 40). The annealing of 5’UAR/c5’UAR, 5’cHP/c5’cHP and 5’UAR/3’UAR were then performed by adding DENV2C at a protein to ODN molar ratio of 2:1, to ensure aggregation-free conditions. Interestingly, DENV2C caused a dramatic increase in all the three annealing reactions, as both 5’UAR/c5’UAR and 5’cHP/c5’cHP annealing were completed much faster as compared to their respective annealing in the absence of the protein (Fig. S3D and S3F). Unlike in the absence of DENV2C, 5’UAR/3’UAR annealing was completed in ~1 hour in the presence of the protein (Fig. 2C and S3E). The kinetic traces could be adequately fitted with equation [1] and kinetic parameters were obtained for DENV2C promoted 5’UAR/c5’UAR, 5’cHP/c5’cHP and 5’UAR/3’UAR annealing (Table 1).

Similar to the absence of DENV2C, the fast component, *k*_obs1_, varied linearly and the slow component, *k*_obs2_, showed hyperbolic dependence with increasing [c5’UAR] during DENV2C promoted 5’UAR/c5’UAR annealing (Fig. 2E and 2H). This suggests a similar reaction mechanism as to that proposed by Scheme 1. Thus, similar to other RNA chaperone promoted ODN annealing (31, 43) and taking acquired kinetic data into account, a reaction mechanism with single kinetic pathway involving a single 5’UAR/DENV2C complex (5’UAR_1_) can be proposed:

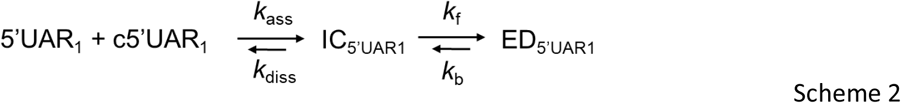

where a fast pre-equilibrium intermediate complex, IC_5’UAR1_, precedes the formation of the final stable extended duplex, ED_5’UAR1_, through a monomolecular reaction (28). The formation of IC_5’UAR1_ is governed by the second order association constant, *k*_ass_, and the first order dissociation constant, *k*_diss_, whereas the formation of ED_5’UAR1_ is governed by the forward interconversion kinetic rate constant, *k*_f_. Again, we validated 5’UAR/c5’UAR annealing mechanism in the presence of DENV2C by using the Dynafit numerical resolution software (Table S1) (30–34).

To gain further insight into the annealing mechanism, we evaluated the temperature dependence of the *k*_obs_ values. The relationship revealed positive enthalpy values of the transition states of 28.6 (± 0.9) kcal/mol and 18.0 (± 1.0) kcal/mol for the fast and slow components, respectively (Fig. 4 and Table 2). These values indicate that 5’UAR/c5’UAR annealing promoted by the DENV2C involves pre-melting of ~5 to ~3 bp in the 5’UAR hairpin structure (35, 36).

Next, the linear and the hyperbolic dependence of *k*_obs1_ and *k*_obs2_, respectively with increasing [c5’cHP] (Fig. S3F and S3H) during the DENV2C promoted 5’cHP/c5’cHP annealing implies that the reaction mechanism follows a single kinetic pathway scheme with an accumulation of IC_cHP1_ followed by its interconversion from ED_cHP1_, similar to the one proposed by scheme 1 (Scheme b in the supplementary). Comparably, DENV2C-promoted 5’UAR/3’UAR annealing also showed linear and hyperbolic dependence of *k*_obs1_ and *k*_obs2_, respectively with increasing [3’UAR] (Fig. 2F and 2I), again indicating a reaction scheme similar to that of scheme 1 (Scheme c in the supplementary).

Comparison of the association constants, *k*_ass_, suggests DENV2C chaperones the rate of intermediate complex formation by ~2 orders of magnitude during both 5’UAR/c5’UAR and 5’cHP/c5’cHP annealing (Table 1). Similarly, in the presence of DENV2C, the rate of intermediate complex conversion to the extended duplex increases by ~5-folds for the 5’UAR/c5’UAR annealing and by ~20-folds for the 5’cHP/c5’cHP annealing (comparing corresponding *k*_f_ values in Table 1). Interestingly, we observed a ~5-fold increase in the *k*_ass_ values of the 5’UAR/3’UAR (39.4 × 10^3^ M^−1^s^−1^) annealing as compared to the *k*_ass_ value of the 5’UAR/c5’UAR (11.9 × 10^4^ M^−1^s^−1^) annealing in the presence of DENV2C. This indicates that the presence of DENV2C probably lowers the kinetic barrier, present due to the non-complementary intermolecular base pairs between 5’UAR and 3’UAR hairpins (Fig. S3B). This result is also in line with the lower enthalpy values obtained for the transition states of 8.2 (± 0.4) kcal/mol for the fast component (Fig. 4 and Table 2) during DENV2C-promoted 5’UAR/3’UAR annealing and implies that the formation of IC_3’UAR1_ requires the pre-melting of only ~2 bp (Scheme c in the supplementary) as compared to the ~5 bp during the formation of IC_5’UAR1_ (Table 1 and Scheme 2). Although IC_3’UAR1_ formed rapidly as compared to IC_5’UAR1_, the ~8-folds increase in its dissociation rate constant, *k*_diss_ showed that the structural stability of such intermediate complex is substantially low, probably due to the intermolecular base pair mismatches in the 5’UAR/3’UAR duplex as compared to the 5’UAR’c5’UAR duplexes (Fig. S3A and S3B).

### DENV2C switches nucleation of 5’UAR/c5’UAR and 5’cHP/c5’cHP annealing through kissing-loop intermediates to stem-stem interactions but not of the 5’UAR/3’UAR annealing

We characterize the molecular mechanisms of 5’UAR/c5’UAR and 5’cHP/c5’cHP annealing by investigating the role of the hairpin loop. To determine whether the annealing was nucleated through kissing-loop intermediates, we used 5’UAR-ULoop and 5’cHP-ULoop mutants, where the G_9_, C_10_, A_11_, G_12_, A_13_ and A_10, 11, 12 and 13_, C_14_ residues, respectively, were changed to U residues (Fig. 1B) in order to decrease the complementarities between the central loops. For both 5’UAR-ULoop/c5’UAR and 5’cHP-ULoop/c5’cHP annealing reactions, no increase in FAM fluorescence was observed (Fig. 5A and Fig. S5A). This drastic decrease in annealing reactions with the loop mutants strongly suggest that both 5’UAR/c5’UAR and 5’cHP/c5’cHP annealing reactions are mainly nucleated through the kissing-loop intermediates. In contrast, in the presence of DENV2C both 5’UAR-ULoop/c5’UAR and 5’cHP-ULoop/c5’cHP kinetic reactions were found similar to that of DENV2C promoted 5’UAR/c5’UAR and DENV2C promoted 5’cHP/c5’cHP annealing (Fig. 5B and Fig. S5B), respectively. No decrease in annealing reaction rates (values provided in Fig. 5 legend) with the loop mutants indicate that the role of hairpin loops in DENV2C promoted 5’UAR/c5’UAR and DENV2C promoted 5’cHP/c5’cHP annealing is limited and hence, both kinetic reactions are prominently nucleated through the stems.

**Figure 5:**
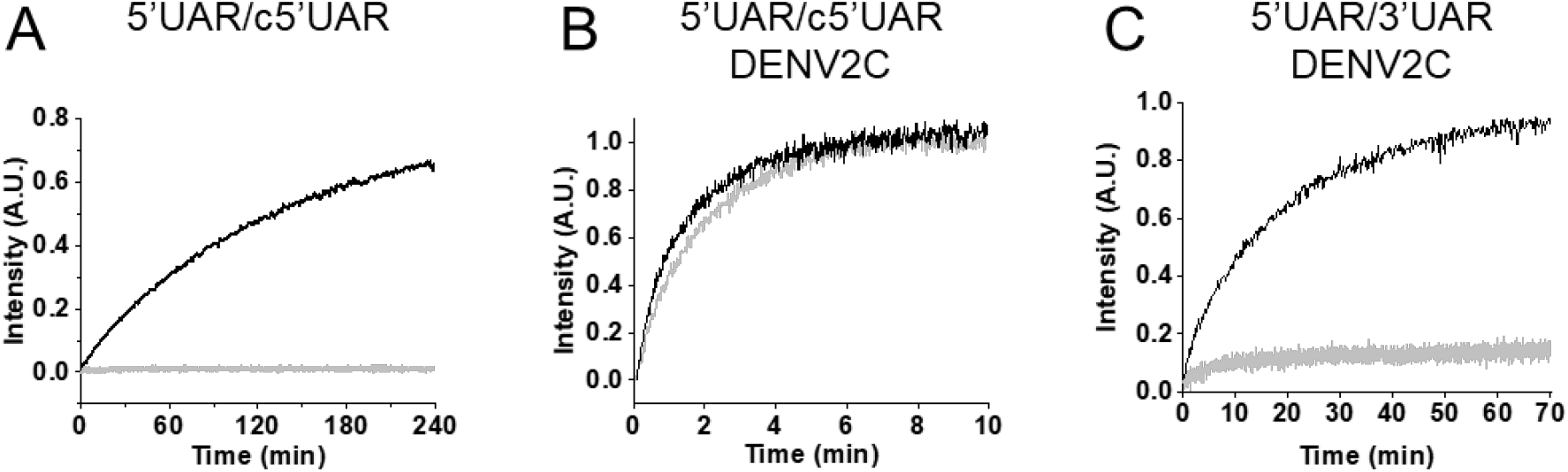
Real-time progress curves the 5’UAR mutants. Progress curve of dual-labeled 5’UAR (black) or 5’UAR-ULoop mutant (grey) with its complementary sequences (A and B) c5’UAR or (C) 3’UAR in the absence and presence of DENV2C, respectively. (A) 10 μM c5’UAR, (B) 700 nM c5’UAR and (C) 1 μM 3’UAR were used for the annealing reaction with either 10 nM of 5’UAR (black traces) or 10 nM of 5’UAR-ULoop (grey traces). (B) Fitting of the progressive curves using Equation [1] provided values for the 5’UAR/c5’UAR reaction in the absence (*k*_obs1_ and *k*_obs2_ are 4.0 × 10^−2^ s^−1^ and 4.4 × 10^−3^ s^−1^) and the presence (*k*_obs1_ and *k*_obs2_ of 4.5 × 10^−2^ s^−1^ and 4.7 × 10^−3^ s^−1^) of DENV2C. Excitation and emission wavelengths used were 480 nm and 520 nm, respectively.

Taken together, these results showed that DENV2C switches the annealing mechanism of complementary sequences during (+)/(−) ds-RNA formation from predominantly through kissing-loop intermediates to propagate through stem-stem interactions. It is possible that the melting of hydrogen bonds in the stem region of the 5’UAR or the c5’cHP hairpin increases due to its interaction with DENV2C protein and this in turn lead to switching of the annealing mechanism. Interestingly, a drastic decrease in the DENV2C promoted 5’UAR-ULoop/3’UAR annealing was observed as compared to its counterpart the DENV2C promoted 5’UAR/3’UAR annealing (Fig. 5C). The result indicates a vital role of the hairpin loop in the DENV2C promoted 5’UAR/3’UAR annealing. Taken together the result states that although DENV2 accelerates the 5’UAR/3’UAR annealing, the role of hairpin loop remains crucial in this annealing. Thus, the annealing mechanism of complementary sequences during the (+) RNA circularization predominantly propagates through loop intermediates.

Since the presence of DENV2C altered the role of hairpin loops in annealing pathways of 5’UAR/c5’UAR and 5’cHP/c5’cHP as compared to their annealing in absence of the protein but not for the 5’UAR/3’UAR annealing, DENV2C probably plays a role in modulating the annealing processes during (+)/(−) ds-RNA formation and the (+) RNA circularization.

### DENV2C protein facilitates 5’UAR/c5’UAR 5’UAR/3’UAR annealing by decreasing the intrinsic dynamics of the 5’UAR hairpin

To further understand how DENV2C chaperones the 5’UAR/c5’UAR and 5’UAR/3’UAR annealing during (+)/(−) ds-RNA formation and (+) RNA circularization respectively, we measured the intrinsic dynamics of the 5’UAR in the presence and in the absence of the protein. We used FRET-FCS (44, 45) which analyses the fluctuations in FRET efficiency, caused by doubly labeled 5’UAR conformational fluctuations. Unlike FCS, FRET-FCS measurers the proximity ratio (*p*), which is function related to FRET efficiency (45–50) and depends on the separation between donor and acceptor but not on the position of the molecules in the observation volume. Thus, the correlation function of the proximity ratio *p* (*Gp*) provides information about structural dynamics of the various 5’UAR conformations. In this case, the 5’UAR might exist in closed, partially-open and fully-open hairpin conformations and thus, the correlation function of *p* can be fitted to a stretched exponential equation (Fig. 6). The equation provides an effective relaxation time (τ_p_), and a stretch parameter (β) describing the heterogeneity of the system. A value of 1 for β indicates that the system displays normal two-state Arrhenius kinetics, with one discrete energy barrier, while a value of 0 for β indicates a continuum of equal energy barriers. The approach was calibrated as mentioned by Sharma KK et al. (49) and effective relaxation time (τ_p_) associated with the motion of the 5’UAR hairpin extremities, were determined. Addition of DENV2C to the 5’UAR hairpin resulted in a ~2-fold increase in values of τ_p_ (from 74 ± 21 μs in the absence to 135 ± 28 μs in the presence of DENV2C) (Fig. 6A), suggesting the hampered fluctuation of 5’UAR hairpin extremities. The β values of 0.21 (± 0.04) in the absence and 0.25 (± 0.06) in the presence of DENV2C showed that the 5’UAR hairpin exists in more than 2 conformations and that the addition of DENV2C does not alter heterogeneity of the system. However, it is possible that one of the 5’UAR hairpin conformations is favored in the presence of the protein.

**Figure 6:**
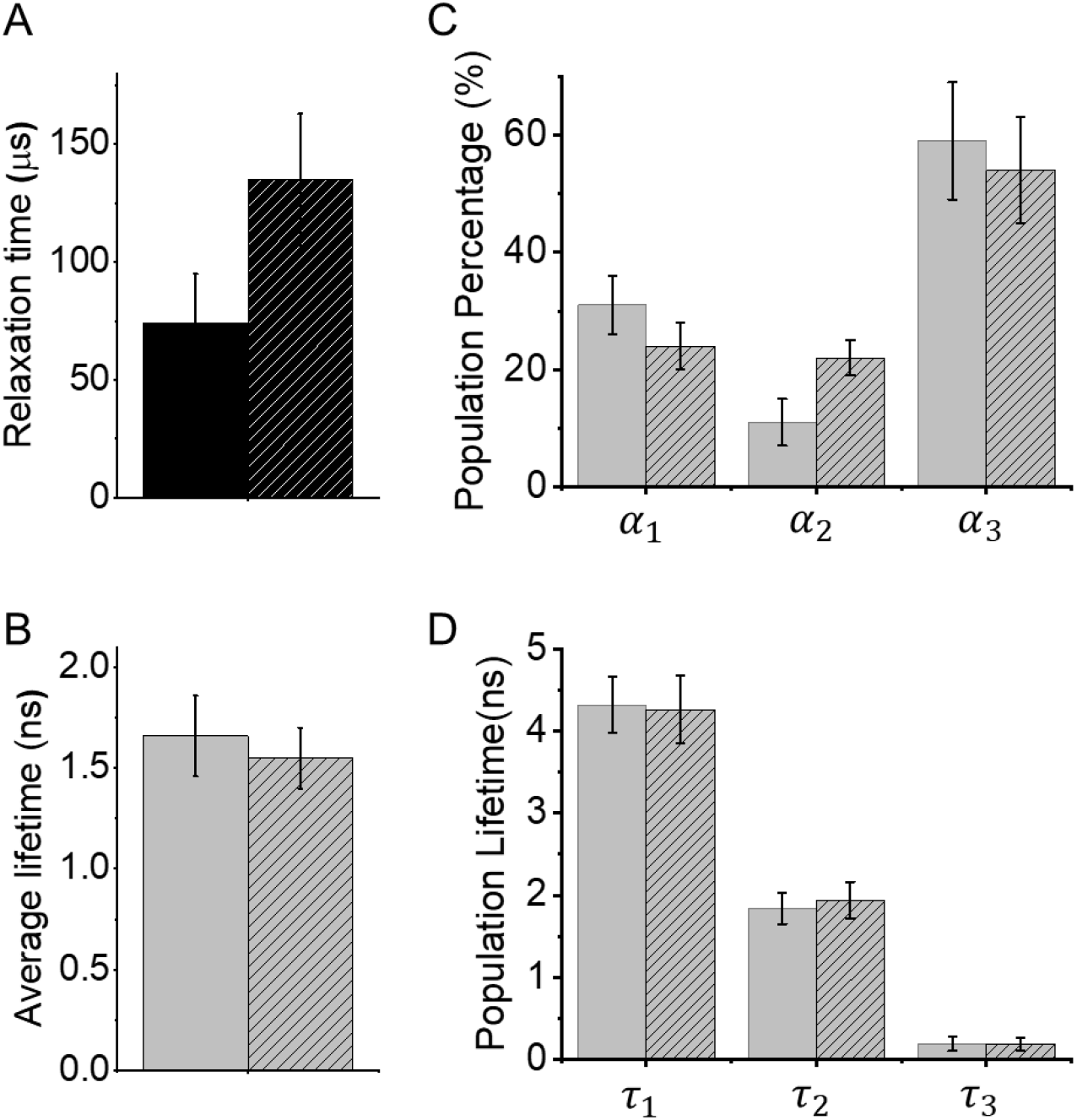
FRET-FCS and trFRET of 5’UAR in the presence and the absence of DENV2C. (A) The relaxation time, τ_*p*_, of 5’UAR in the absence (solid) and presence (striped) of DENV2C. An autocorrelation function of proximity ratio, *G_p_ (τ)*, was constructed using: 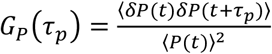 and fitted to a stretched exponential equation (47):

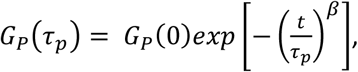

where τ_*p*_ corresponds to the effective relaxation time associated with the correlated motion and β is a stretch parameter. (B) The lifetime traces for doubly labeled 5’UAR were fitted to tri-exponential decay model in the average fluorescence lifetimes (<τ_f_>) of FAM for donor was calculated in the (solid) absence and in the (striped) presence of DENV2C. Fitting of the fluorescence decay provided three discrete lifetimes, τ_1_, τ_2_ and τ_3_ (C and D) with the corresponding fraction of each population as α_1_, α_2_ and α_3_, respectively in the (solid) absence and (striped) the presence of DENV2C added at protein:ODN ratio of 2:1. Error bars represents standard deviations of at least three repeats.

We investigated the notion of the favored 5’UAR conformation among its various conformations using trFRET. In trFRET, the energy transfer from donor to acceptor influences the donor fluorescence lifetime, τ, in a distant dependent manner. The closer the donor and acceptor dyes are, the faster the donor dye relaxes into ground state and the shorter the lifetime. The fluorescence decay of 5’UAR had an average lifetime of 1.7 ± 0.2 ns (Fig. 6B) and was best fitted with three discrete lifetime components of ~4.3 ns (<τ_1_>), ~1.9 ns (<τ_2_>) and ~0.19 ns (<τ_3_>) having populations of 31 ± 5% (α_1_), 11 ±4% (α_2_) and 59 ± 10% (α_3_), respectively (Fig. 6C and 6D). Note that we cannot exclude the existence of conformations with very short lifetimes or with lifetimes close to the three principal components. Those would not be resolvable within our experimental conditions. We refer to the fluorescence lifetime of closed, partially-open and fully-open conformations of the 5’UAR hairpins as <τ_3_>, <τ_2_> and <τ_1_> and their corresponding populations as α_3_, α_2_ and α_1_.

No or marginal difference in the average lifetime of the donor was observed after addition of DENV2C to the 5’UAR hairpin. However, a ~2-fold increase in the value of α_2_ was observed that likely happened at the expense of both α_1_ and α_3_ populations (Fig. 6C). The result indicates that as an RNA chaperone with the ability to both dissociate and anneal RNA (20), DENV2C could be effectively unwinding high FRET populations (<τ_1_> and α_1_) and annealing low FRET populations (<τ_3_> and α_3_) to allow the 5’UAR hairpin to reach its most favored kinetic conformation (<τ_2_> and α_2_). Overall, the results from FRET-FCS and trFRET explain that DENV2C probably exerts its RNA chaperone activities on the 5’UAR by modulating its intrinsic dynamics as well as by decreasing kinetically trapped unfavorable conformations. Such mechanistic behavior of the DENV2C is in line with the ‘entropy exchange model’ in which a highly flexible protein, like DENV2C, undergoes disorder-to-order transition upon binding to RNA that in turn leads to the melting of the RNA structure through an entropy exchange process (23).

## DISCUSSION

Large-scale genome rearrangements, such as the (+) RNA circularization and the (−) RNA synthesis are essential during viral RNA replication in the dengue virus life cycle. RNA chaperones such as DENV2C are vital for preventing RNA misfolding by allowing RNA to escape unfavorable kinetic traps. In this study, we characterized the role of DENV2C as an RNA chaperone by investigating the annealing kinetics of the 5’UAR and the 5’cHP to their corresponding complementary sequences during (+)/(−) ds-RNA formation and the (+) RNA circularization. The hybridization of 5’UAR and 5’cHP sequences are important tertiary interactions during DENV replication and essential for its genome amplification (7). A systemic monitoring of real-time annealing kinetics of 5’UAR and 5’cHP mutants in the presence and the absence of DENV2C were performed. It was observed that annealing reaction during (+)/(−) ds-RNA formation propagates through the kissing-loop intermediates albeit rather slowly, taking several hours to reach completion (Fig. 2A and Fig. S3A). This slow annealing kinetics can be related with the requirement of the complementary sequences to be in a reactive conformation and proper orientation in order to nucleate the intermediate complex. Formation of the IC_5’UAR_ intermediate involves melting of the G_7_-C_15_ and A_8_-U_14_ base pairs near the bottom of the 5’UAR loop. The resulting intermediate complex, IC_5’UAR_, could be stabilized by up to 9 intermolecular base pairs (Fig 7A). The IC_5’UAR_ is then converted into the ED_5’UAR_ in a rate limiting step associated with the transition enthalpy of ~18 kcal mol^−1^, that likely corresponds to the melting of the three next A_6_-U_16_, G_5_-C_17_ and A_4_-U_18_ base pairs in the middle region of the 5’UAR stem. This melting is probably the bottleneck for the interconversion into ED_5’UAR_ (Fig. 7A).

**Figure 7:**
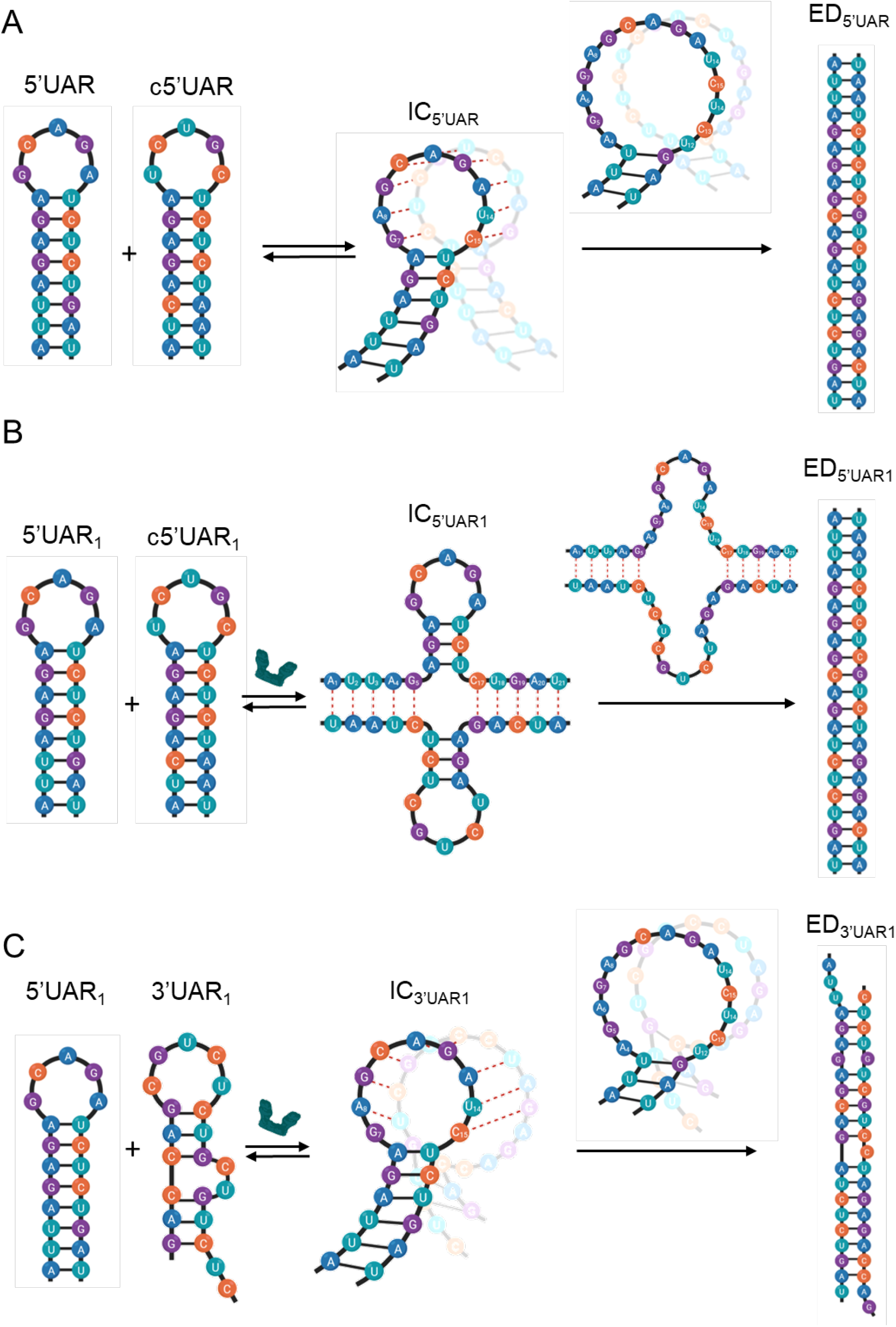
Proposed reaction mechanisms for annealing of 5’UAR/c5’UAR and 5’UAR/3’UAR in presence and absence of DENV2C. (A) Proposed mechanism for 5’UAR/c5’UAR involves the annealing kinetics showing an accumulation of an intermediate complex (IC_5’UAR_) and a one-pathway reaction scheme. Annealing is nucleated via kissing-loop intermediate (IC_5’UAR_), consisting of hydrogen bonding between complementary nucleotides in the loops of the 5’UAR and the c5’UAR (red dashed line). Formation of the IC_5’UAR_ involves melting of two (G_7_-C_15_ and A_8_-U_14_) base pairs near the bottom of the 5’UAR loop. After which, interconversion from IC_5’UAR_ to ED_5’UAR_ takes place, likely involving the melting of the three next (A_6_-U_16_, G_5_-C_17_ and A_4_-U_18_) base pairs in the middle region of the 5’UAR stem. (B) The DENV2C promoted 5’UAR/c5’UAR annealing starts with opening of the 5’UAR hairpin in the stem region, allowing annealing via the stems. Formation of the IC_5’UAR1_ involves melting of the five (A_1_-U_21_, U_2_-A_20_, U_3_-G_19_, A_4_-U_18_ and G_5_-C_17_) base pairs in the lower 5’UAR stem. Interconversion from IC_5’UAR1_ to ED_5’UAR1_ involves melting of the three remaining (A_6_-U_16_, G_7_-C_15_ and A_8_-U_14_) base pairs in the upper part of 5’UAR stem. (C) The DENV2C promoted 5’UAR/3’UAR annealing takes place predominantly through the kissing-loop intermediates, with IC_3’UAR1_ formation likely involving the melting of the two (G_7_-C_15_ and A_8_-U_14_) base pairs near the bottom of the 5’UAR loop. Interconversion from IC to ED likely involves the melting of the three remaining (A_6_-U_16_, G_5_-C_17_ and A_4_-U_18_) base pairs in the middle region of the 5’UAR stem.

Interestingly, in the presence of DENV2C, the annealing mechanism of 5’UAR/c5’UAR switched from mainly nucleating through the kissing-loop intermediates and proceeded via the stem-stem interactions. Formation of the 5’UAR/c5’UAR intermediates in the presence of DENV2C probably involves melting of the A_1_-U_21_, U_2_-A_20_, U_3_-G_19_, A_4_-U_18_ and G_5_-C_17_ base pairs in the lower 5’UAR stem. The resulting intermediate complex (IC_5’UAR1_) could be stabilized by up to 10 intermolecular base pairs (Fig 7B). The IC_5’UAR1_ is further converted into the ED_5’UAR1_, which most probably relies on the conformational rearrangement and melting of the three remaining A_6_-U_16_, G_7_-C_15_ and A_8_-U_14_ base pairs in the unstable region of the upper 5’UAR stem. In this respect, the ~5-fold increased interconversion rate of IC_5’UAR1_ probably results from its stability due to larger number of intermolecular base pairs and/or more favorable conformation for the subsequent conversion into the ED_5’UAR1_.

In contrast, a vital role of the loop in the DENV2C-promoted 5’UAR/3’UAR annealing forced the reaction mechanism to proceed through the kissing-loop intermediate, IC_3’UAR1_. Thus, similar to the 5’UAR/c5’UAR intermediates in the absence of the protein, the formation of 5’UAR/3’UAR intermediates involves melting of the G_7_-C_15_ and A_8_-U_14_ base pairs in the upper region of the 5’UAR stem and the resulting intermediate complex could be stabilized by up to 8 intermolecular base pairs (Fig. S3B) followed by its interconversion to the extended duplex by the melting of the three next A_6_-U_16_, G_5_-C_17_ and A_4_-U_18_ base pairs in the middle region of the 5’UAR stem (Fig 7C). The initial three bp at 5’ end of the 5’UAR stem are non-complementary to the corresponding nucleotides in the 3’UAR stems (Fig. S3B). This may hinder the propagation of the DENV2C promoted 5’UAR/3’UAR annealing through the hairpin stems. In addition, the lower transition enthalpy of ~8 kcal mol^−1^ indicated a rapid formation of IC_3’UAR1_ and in turn a ~5-fold increased association constant compared to the DENV2C promoted 5’UAR/c5’UAR annealing (Table 1). However, IC_3’UAR1_ is probably stabilized with only ~5 intermolecular base pairs (involving A_8_, G_9_, C_10_, A_11_ and G_12_ of 5’UAR hairpin) which lead to its ~8-fold increased dissociation rate constant compared to the DENV2 promoted 5’UAR/c5’UAR annealing (Table 1). Apart from reduced intermolecular base pairing, the inferior stability of IC_3’UAR1_ could also be attributed to non-complementarities at the bottom of the 5’UAR loop involving G_7 and 12_ (Fig. S3B) and melting of three base pairs in the stable 5’UAR stem. Therefore, the combination of mismatches defining the stability of the ODN duplexes and the structured RNA elements attacked by DENV2C during ODN annealing (either hairpin loop or hairpin stem) probably dictates the processes of (+)/(−) ds-RNA formation and the (+) RNA circularization.

Similar to other RNA chaperones (19, 51–56), FRET-FCS and trFRET data showed that DENV2C also functions by binding to RNA and affecting their intrinsic dynamics, allowing them to escape their thermodynamically stable, but non-functional state and explore other states until the RNA is finally able to find its functionally correct state (57). A comparison between different RNA chaperones suggests that DENV2C, HIV-1 NCp7 and hepatitis C virus (HCV) core protein accelerates annealing of complementary ODNs up to 2 to 3 orders of magnitude (30, 40, 58), indicating similar chaperoning potential for the three proteins. However, DENV2C accelerates annealing of complementary ODNs at two equivalent of protein molecule while HIV-1 NCp7 and HCV core require ~six and ~two equivalent of protein molecules, respectively, to show similar chaperone potential. This suggests that one molecule of either DENV2C or HCV core could be as active as three molecules of HIV-1 NCp7. Interestingly, both DENV2C and HCV core are existing as homodimer while HIV-1 NCp7 is a monomeric protein *in vitro* (59, 60). Thus, like HCV core, the superior chaperone activity of DENV2C compared to HIV-1 NCp7 is likely a consequence of the stronger ‘nucleic acid aggregating’ properties, in line with the efficient oligonucleotide aggregation observed by FCS (Fig. S4). This propensity of the DENV2C to neutralize the negatively charged nucleic acids and to promote their aggregation is probably related to the highly flexible and unstructured helix 1 (22, 59, 61) as compared to the folded HIV-1 NCp7 structure with two zinc fingers (60, 62). Moreover, HIV-1 NCp7 promotes ODN annealing through kissing loop intermediates (28) while the HCV core mainly chaperone the annealing through the stems (30), the DENV2C can propagate annealing either through kissing-loop intermediates (during the (+) RNA circularization) or through the stems (during (+)/(−) ds-RNA formation). Taken together, our results showed that the DENV2C have stronger nucleic acid annealing activity at low protein/ODN ratios when compared to HIV-1 NCp7, while it can modulate annealing pathways when compared to HCV core. Despite of structural differences between these three proteins, the conserved nucleic acid chaperone properties suggest that they are required in different viruses, in line with the conservation of RNA chaperoning in *Flaviviridae* core proteins (21). Such properties are not observed in other DENV proteins. Other viral proteins which interact with genomic RNA are NS5 polymerase and NS3 helicase. However, the ribozyme activity assay showed that both NS5 polymerase and NS3 helicase have no or limited RNA chaperoning activity (20). Thus, DENV2C is likely to be the only viral protein responsible for preventing misfolding of RNA during replication. In addition, our results suggest that the DENV2C plays a probable role in modulating (+)/(−) ds-RNA formation and the (+) RNA circularization. These abilities of DENV2C may be critical for genomic RNA dimerization in DENV replication and RNA packaging, as well in facilitating recombination between various DENV genotypes and subtypes to increase viral variability.

