## Supplementary material for "Dengue virus strain 2 capsid protein switches the annealing pathway and reduces the intrinsic dynamics of the conserved 5’ untranslated region"

**Dynafit simulation procedure**

Simulated progress curves were obtained at different concentration of c5'UAR (1 µM, 3 µM, 5 µM, 8 µM, 10 µM, 15 µM, 20 µM, 25 µM, 30 µM, 40 µM and 50 µM) using elementary rate constants, $k_{obs1}$, $k_{obs2}$ and $k_f$, from Table 1 and according to the following Scheme (1–3):

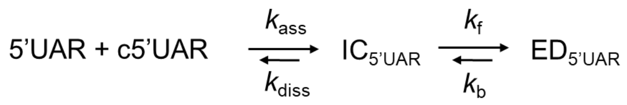


To obtain simulated progress curves, the concentration of 5'UAR was kept at a constant value of 10 nM. Progress curves of the 5'UAR/c5'UAR reaction were fitted to Equation [1] to obtain values for $k_{obs1}$ and $k_{obs2}$ with increasing concentration of c5'UAR. The fast and slow reaction rates, $k_{obs1}$ and $k_{obs2}$, were plotted against the concentration of c5'UAR (Fig. S1A and Fig. S1B) and fitted using Equation [2] and [3], respectively. The generated kinetic parameters are shown in Table S1.

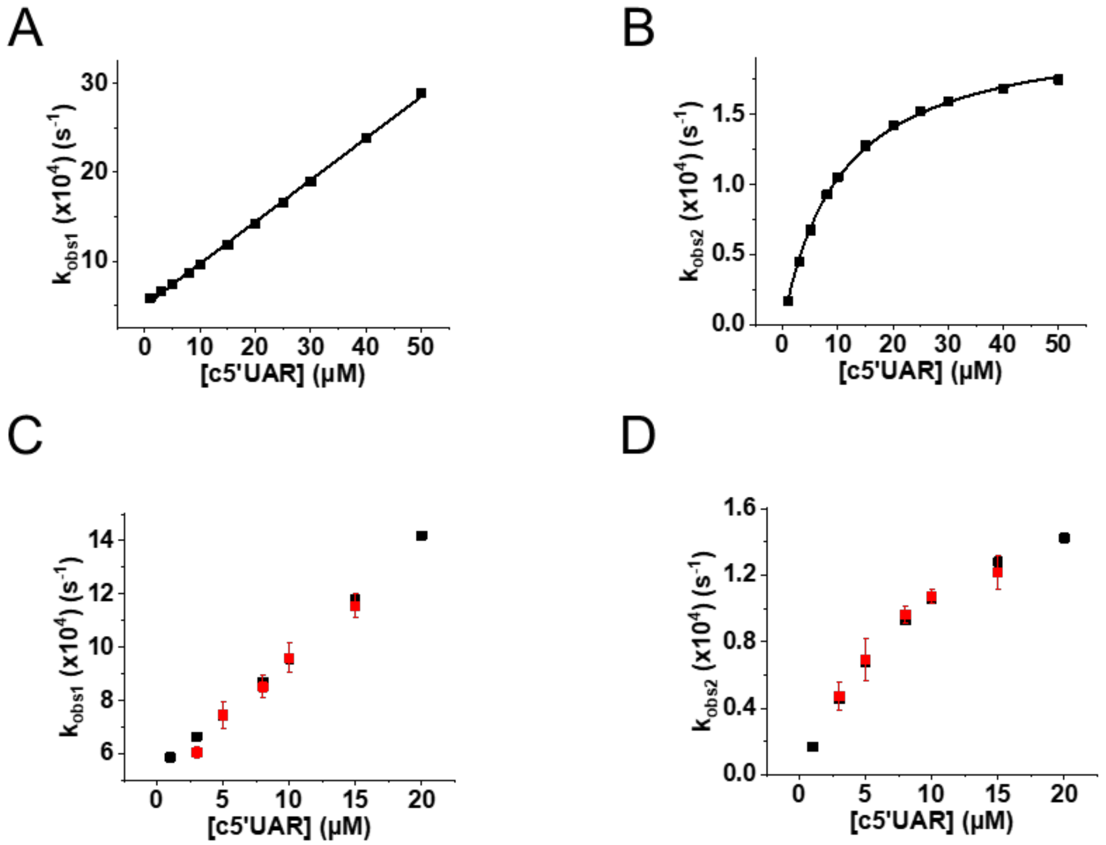


**Figure S1 – Simulated kinetic parameters of 5'UAR/c5'UAR annealing.** The fast ($k_{obs1}$) (A) and slow ($k_{obs2}$) (B) components for 5'UAR/c5'UAR annealing were determined by fitting simulated progress traces, which were obtained by using empirical kinetic parameters in Table 1 and increasing concentrations of c5'UAR. The solid lines correspond to the fit of the fast component (A) with Equation [2] and slow component (B) with Equation [3]. The values are provided in Table S1. Comparison between kinetic parameters obtained for fast (C) and slow (D) components using experimental (red squares) and simulated (black squares) approach. The similar values of simulated kinetic parameters to that of experimental kinetic parameter values supports the proposed reaction Schemes in the article.

**Table S1 – Kinetic parameters of 5'UAR/c5'UAR, 5'cHP/c5'cHP and 5'UAR/3'UAR annealing and their mutants in the absence and presence of DENV2C obtained by fitting simulated progression curves.**

| Doubly labeled ODN | Complementary ODN | DENV2C/ODN ratio | $k_{ass1}$ ($M^{-1}s^{-1}$) × $10^{-3}$ | $k_{diss1}$ ($s^{-1}$) × $10^{4}$ | $K_a$ ($M^{-1}$) × $10^{-5}$ | $k_f$ ($s^{-1}$) × $10^{4}$ |
|---|---|---|---|---|---|---|
| 5'UAR | c5'UAR | 0 | 0.047 (±0.005) | 5 (± 0.1) | 0.96 (± 0.03) | 2.1 (± 0.02) |
| 5'UAR | c5'UAR | 2 | 8.3 (± 0.02) | 39 (± 0.05) | 20.4 (± 0.4) | 9.7 (± 0.03) |
| 5'UAR | 3'UAR | 2 | 39.2 (± 0.02) | 260 (± 0.5) | 14.5 (± 0.1) | 13.6 (± 0.03) |
| 5'cHP | c5'cHP | 0 | 0.084 (±0.001) | 2.8 (± 0.03) | 2.9 (± 0.03) | 0.51 (± 0.01) |
| 5'cHP | c5'cHP | 2 | 7.3 (± 0.01) | 73 (± 0.05) | 9.8 (± 0.1) | 10.6 (± 0.03) |

Kinetic rate constants were calculated from the dependence of the $k_{obs}$ values on the concentration of the unlabeled complementary ODN, as indicated in Figure S1. The $k_{ass}$ and $k_{diss}$ values were calculated using Equation [2], while the $K_a$ and $k_f$ values were calculated using Equation [3]. Simulated kinetic parameters and empirical values (Table 1) were similar, supporting the proposed reaction Schemes in the article.

**Supplementary Schemes**

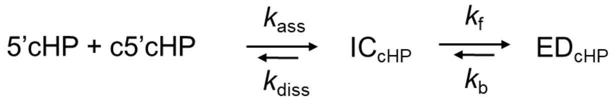


**Scheme a**

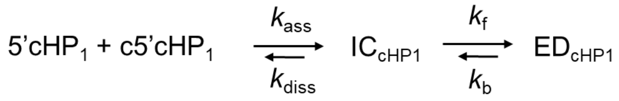


**Scheme b**

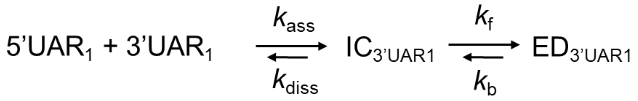


**Scheme c**

According to the above-mentioned reaction schemes, a reaction mechanism with a single kinetic pathway involving a single initiation complex (ODN alone, 5'cHP, or in complex with DENV2C, 5'cHP$_1$ and 5'UAR$_1$) is proposed. In the reactions, a fast pre-equilibrium intermediate complex, $IC_{cHP}$, $IC_{cHP1}$ and $IC_{3'UAR1}$, precedes the formation of the final stable extended duplexes, $ED_{cHP}$, $ED_{cHP1}$ and $ED_{3'UAR1}$, through a monomolecular reaction. The formation of ICs are governed by the second order association constants, $k_{ass}$, and the first order dissociation constants, $k_{diss}$, whereas the formation of EDs are governed by the forward interconversion kinetic rate constants, $k_f$.

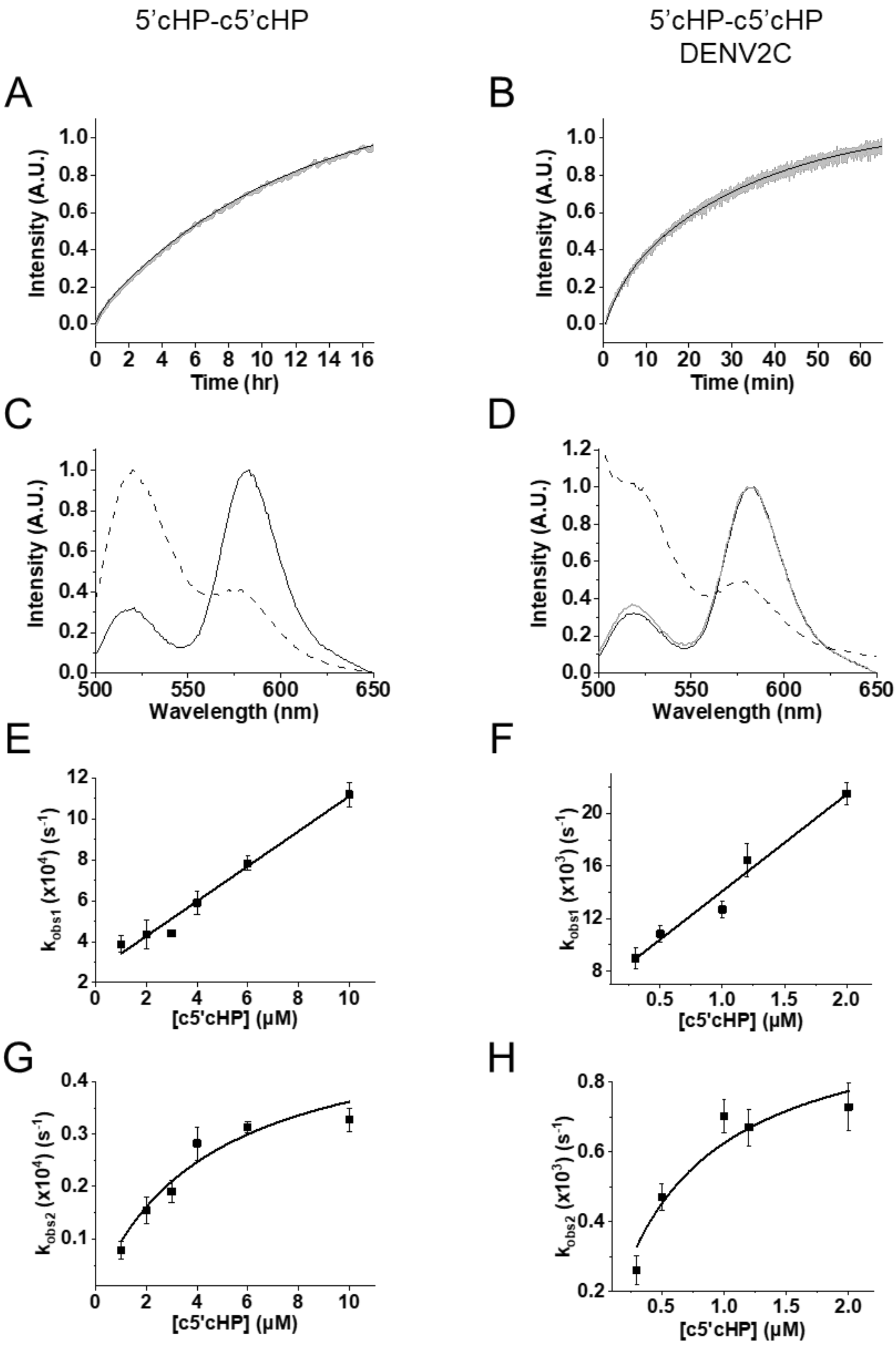


**Figure S2 – Real-time progress intensity curves (A-B), emission spectra (C-D) and kinetic parameters (E-H) of 5'cHP/c5'cHP annealing in the absence (A, C, E and G) and presence (B, D, F and H) of DENV2C.** (A and B) Real-time intensity traces (grey) and their fits to Equation [1] (black) to obtain fast ($k_{obs1}$) and slow ($k_{obs2}$) kinetic parameters. Real-time intensity traces of (A) 10 nM 5'cHP with 10 μM c5'cHP and (B) 10 nM 5'cHP with 1 μM c5'cHP in the presence of DENV2C in a protein:ODN molar ratio of 2:1. (C and D) Emission spectra of 5'cHP before (solid black) and after annealing (dotted black) to c5'cHP, at excitation wavelength 480 nm. The solid grey line shows the emission spectra of 5'cHP after addition of DENV2C in a protein:ODN molar ratio of 2:1. $k_{obs1}$ (E and F) and $k_{obs2}$ (G and H) parameters were plotted against the concentration of c5'cHP and fitted to Equation [2] and [3], respectively.

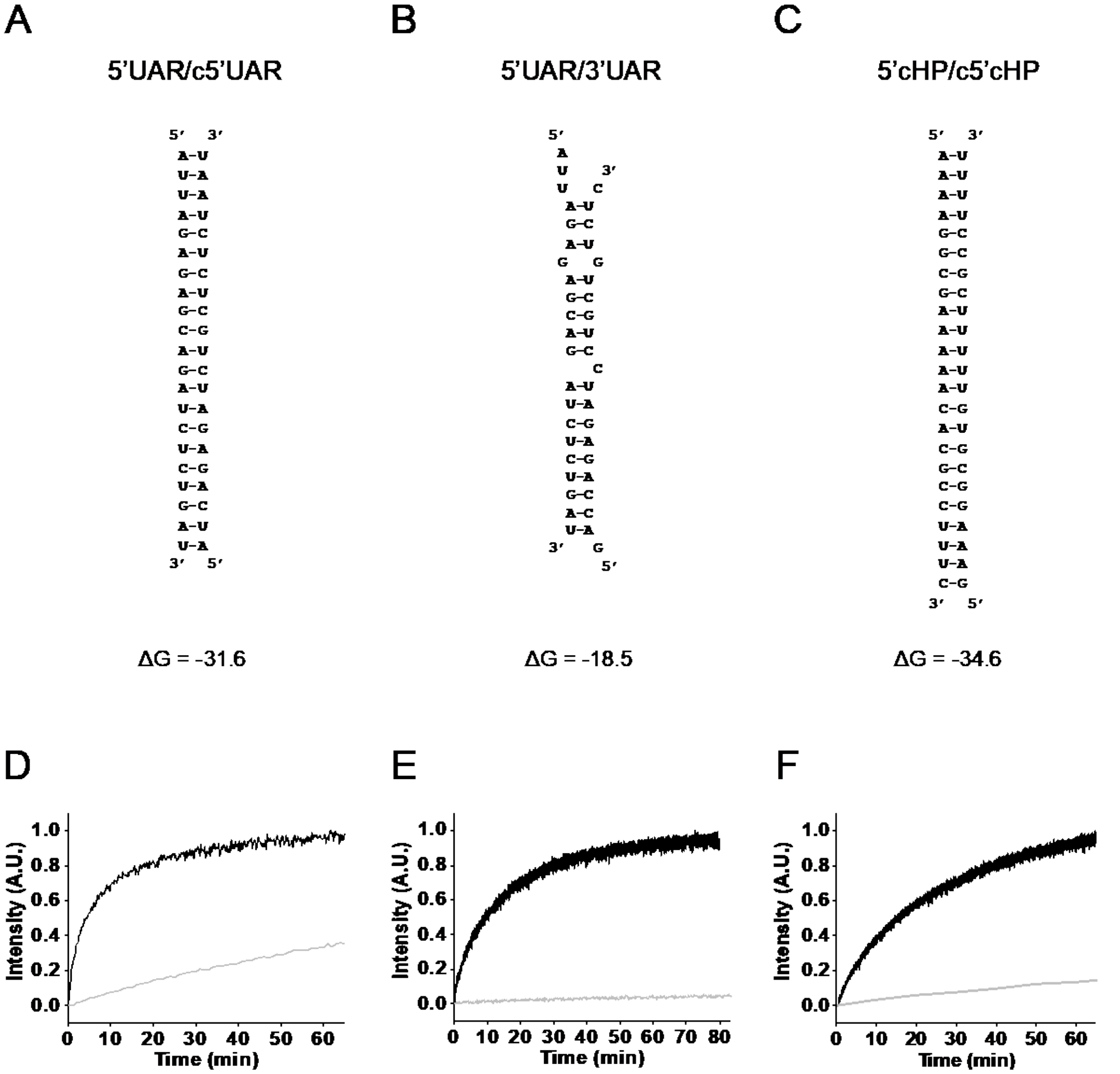


**Figure S3 – Extended duplex structures and comparison of annealing kinetics in the absence and presence of DENV2C.** Extended duplexes (ED) of (A) 5'UAR/c5'UAR, (B) 5'UAR/3'UAR and (C) 5'cHP/c5'cHP. A dash indicates complementary base pairing. The strand on the left is the doubly labelled strand: 5'UAR or 5'cHP. Below each duplex is the Gibbs free energy, ΔG associated with the hybridization of the duplex in kcal/mol. This is determined using the Two-state melting (hybridization) webtool from http://unafold.rna.albany.edu/. Comparison between (D) 5'UAR/c5'UAR, (E) 5'UAR/3'UAR and (F) 5'cHP/c5'cHP reactions in the absence (light grey) and presence (black) of DENV2C. (D) Intensity traces of annealing between 10 nM 5'UAR and 8 μM c5'UAR (light grey), and 1 μM c5'UAR in the presence of DENV2C (black). (E) Intensity traces of annealing between 10 nM 5'UAR and 5 μM 3'UAR (light grey), and 3 μM 3'UAR in the presence of DENV2C (black). (F) Intensity traces of annealing between 10 nM 5'cHP and 10 μM c5'cHP (light grey), and 1 μM c5'cHP in the presence of DENV2C (black).

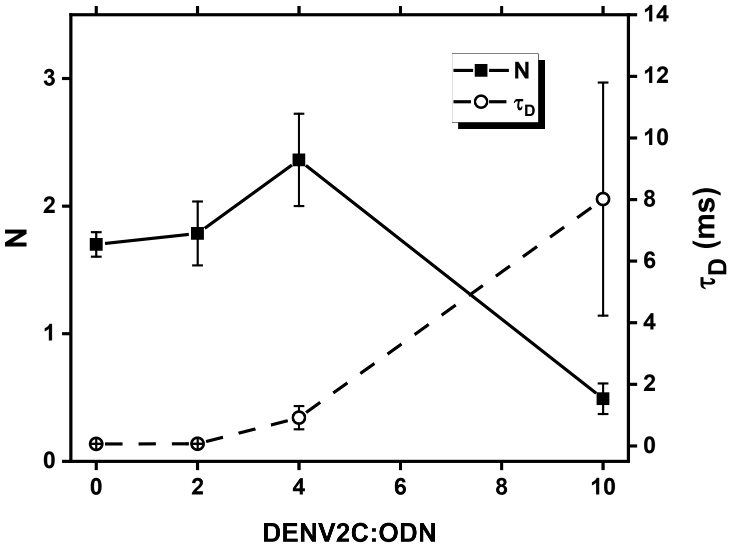


**Figure S4 – The optimal DENV2C:ODN molar ratio to use for annealing reactions is 2:1.** Fluorescence correlation spectroscopy (FCS) was used to monitor aggregation of 10 nM 5’UAR with increasing DENV2C concentrations. 5’UAR diffusion time, $\tau_D$ (circles; dotted line) and number of particles, N (squares; solid line) were measured. A 543 nm continuous wave laser was used, with a 600/50 bandpass filter to monitor fluorescence from 3’TAMRA on 5’UAR. Error bars show standard deviation from three measurements.

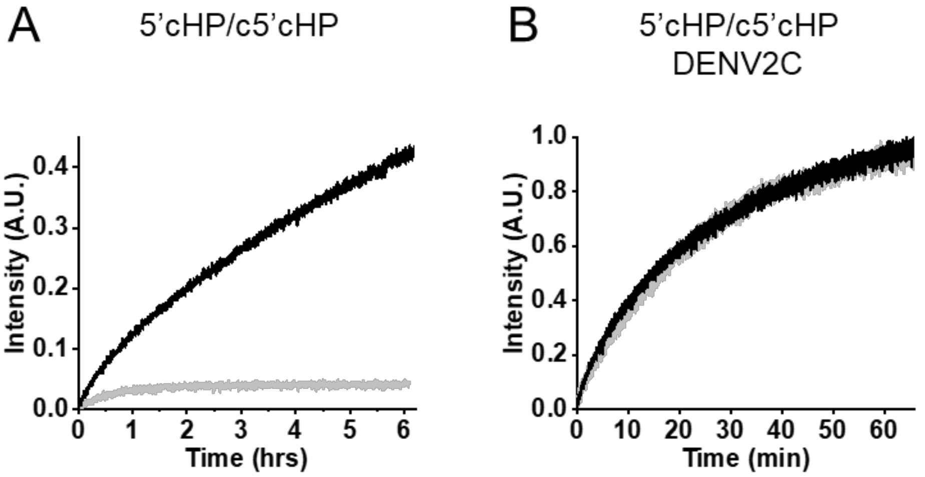


**Figure S5 – Real-time progress intensity traces of dual-labeled 5’cHP (black) or 5’cHP-ULoop mutant (grey) with its complementary sequence c5’cHP in the (A) absence and (B) presence of DENV2C.** (A) 5 μM c5’cHP and (B) 1 μM c5’cHP were used for the reaction with 5’cHP and 5’cHP-ULoop, respectively. (B) Fitting of the real-time intensity curves using Equation [1] provided observed kinetic values for the 5'cHP/c5'cHP reaction ($k_{obs1}$ and $k_{obs2}$ are $1.3 \times 10^{-2}$ $s^{-1}$ and $7.0 \times 10^{-4}$ $s^{-1}$) and the 5'cHP-ULoop/c5'cHP reaction ($k_{obs1}$ and $k_{obs2}$ of $1.4 \times 10^{-2}$ $s^{-1}$ and $6.8 \times 10^{-4}$ $s^{-1}$). Excitation and emission wavelengths used were 480 nm and 520 nm respectively. For reactions performed in the presence of DENV2C, a DENV2C:ODN molar ratio of 2:1 was used.
